# VGLL1 drives therapy resistance in estrogen receptor positive breast cancer

**DOI:** 10.1101/2020.11.29.402842

**Authors:** Carolina Gemma, Chun-Fui Lai, Anup K Singh, Antonino Belfiore, Neil Portman, Heloisa Z Milioli, Manikandan Periyasamy, Sara Abuelmaaty, Alyssa J. Nicholls, Claire M Davies, Naina R. Patel, Georgia M. Simmons, Hailing Fan, Van T M Nguyen, Luca Magnani, Emad Rakha, Lesley-Ann Martin, Elgene Lim, R. Charles Coombes, Giancarlo Pruneri, Lakjaya Buluwela, Simak Ali

## Abstract

Resistance to endocrine therapies (ET) is common in estrogen receptor (ER) positive breast cancer and most relapsed patients die with ET-resistant disease. While genetic mutations provide explanations for some relapsed patients^1^, mechanisms of resistance remain undefined in many cases. Drug-induced epigenetic reprogramming provides possible routes to resistance^2^. By analysing histone H3 lysine 27 acetylation (H3K27ac) profiles in models of ET resistance, we discovered that selective ER down-regulators (SERDs) such as fulvestrant promote epigenetic activation of *VGLL1*, a co-activator for TEAD transcription factors. We show that *VGLL1*, acting via TEADs, promotes expression of genes that drive growth of fulvestrant-resistant breast cancer cells. Pharmacological disruption of VGLL1/TEAD4 interaction inhibits VGLL1/TEAD transcriptional programmes to block growth of the resistant cells and prevents growth. Among the VGLL1/TEAD-regulated genes, we identify *EGFR*, whereby VGLL1-directed EGFR upregulation sensitises fulvestrant-resistant breast cancer cells to EGFR inhibitors. Taken together, our findings identify VGLL1 as a transcriptional driver in ET resistance and advance new therapeutic possibilities for relapsed ER+ breast cancer patients.

## Introduction

ER is the key transcriptional driver of tumour growth in three-quarters of breast cancers patients^3^. As such, ER targeting drugs are effective therapies for most ER+ patients. Despite the success of these treatments, many women develop resistance to these drugs^3–6^, necessitating identification of resistance mechanisms and development of new therapies.

Cell identity is established through epigenetic activation of distal and proximal regulatory elements that direct cell-type-specific gene expression programmes in development and differentiation^7,8^. Cancer cells are also characterised by transcriptional programmes that are frequently defined by cancer type-specific epigenetic states^9–11^, while altered epigenetic landscapes are likely to signpost therapy resistance pathways^12^. Indeed, profiling isogenic ER+ breast cancer cell models of resistance to different ETs for the active transcription histone mark H3K27ac reveals extensive epigenetic reprogramming resulting in sweeping reorganization of enhancer landscapes, findings confirmed with other gene promoter and enhancer mapping approaches^13–15^.

Different classes of ET drugs inhibit ER signalling through distinct mechanisms. Aromatase inhibitors (AI) block estrogen biosynthesis to prevent ER activation and recruitment to DNA^6^. Selective ER modulators (SERMs) like tamoxifen^4^ bind to ER to inhibit its activity, while allowing its recruitment to DNA. Fulvestrant is the prototype clinical drug for ER degraders^16^ that reduce ER levels. Fulvestrant is approved for advanced ER+ breast cancer treatment following relapse on prior ET, either alone or in combination with drugs targeting other pathways, such as CDK4/6 inhibitors^17^. Establishing the clinical value of fulvestrant has galvanised the development of new SERDs with improved pharmacological profiles, several of which have progressed to advanced clinical trials^16,18,19^.

The TEA domain (TEAD) family of transcription factors are implicated in cancer as the DNA binding partners for Yes-associated protein 1 (YAP), and its paralog, transcriptional co-activator with PDZ-binding motif (TAZ)^20–23^. YAP/TAZ are downstream effectors of the Hippo pathway and are required for organogenesis, tissue homeostasis and are important players in cancer initiation and progression, including resistance to cancer therapies^24,25^. By contrast, little is known about the vestigial-like (VGLL) TEAD co-factors. *Vestigial* (vg) is a master regulator of wing development in *Drosophila*, regulating gene expression upon dimerization with *scalloped* (sd), the *Drosophila* homologue of mammalian TEAD^26–29^. Human VGLL1 can substitute for *Drosophila* vg in wing formation^30^, underscoring an evolutionarily conserved role for VGLL1 in TEAD-directed gene expression. Moreover, VGLL1 competes with YAP for binding to TEAD4 *in vitro*, and the VGLL1-TEAD and YAP-TEAD complexes share structural similarities^31,32^.

As the first-in-class clinical SERD, understanding fulvestrant resistance mechanisms is necessary for continued use and the appropriate clinical introduction of newly developed SERDs. Towards this end, we examined H3K27ac profiles in fulvestrant-resistant breast cancer cells and found loss of ER signalling pathways and reprogramming of TEAD-directed gene expression driven by induction of VGLL1, thereby identifying new approaches for treatment of patients who progress on SERDs.

## Results

### VGLL1 induction and downregulation of ER signalling in fulvestrant resistance

Our previous work profiling global H3K27ac patterns revealed distinct epigenetic landscapes in isogenic breast cancer cells resistant to different ETs^13^. As presence of H3K27ac flags active gene promoters^33,34^, we ranked genes according to the ratio of H3K27ac signal in MCF7 cells with acquired resistance to fulvestrant (FULVR)^13^ relative to isogenic fulvestrant-sensitive MCF7 cells. H3K27ac was reduced at ER target genes in FULVR cells, as expected from the ER inhibitory action of fulvestrant (Fig. 1a, b). *VGLL1* (rank=3) and *VGLL3* (rank=5) were among the genes with the greatest increase in H3K27ac signal. Functional annotation analysis^35^ showed that the HIPPO pathway is the most enriched signalling pathway (fold enrichment=4.7, FDR=7.9×10^-6^; Supplementary table 1) in the H3K27ac-enhanced genes (576 genes with log2 fold change (FC) >2). Consistent with the enhancer/promoter reorganisation implied by H3K27ac mapping, RNA-seq gene expression data revealed substantial increases in VGLL1 and VGLL3 expression in FULVR cells (Fig. 1c). Concordant with the H3K27ac results, gene ontology (GO) analysis of the RNA-seq data demonstrated downregulation of estrogen response pathways in FULVR cells (Extended Data Fig. 1a). Immunoblotting and RT-qPCR confirmed that VGLL1 and VGLL3 expression is extremely low in MCF7 cells, indicating that epigenetic reprograming accompanying development of fulvestrant resistance leads to their induction (Fig. 1d, Extended Data Fig. 1b). In MCF7-FULVR cells, YAP and TAZ mRNA levels were substantially lower than those of VGLL1 and VGLL3 and their protein levels were not markedly different in FULVR and MCF7 cells. Interestingly, expression of three of the four TEAD genes was also increased in FULVR cells. Elevated expression of VGLL1 was also clear in other fulvestrant-resistant ER+ cell lines (Extended Data Fig. 1c, d). However, VGLL3 was not commonly upregulated, as was the case for MCF7-FULVR cells, where YAP and TAZ expression was not increased.

**Fig. 1.**
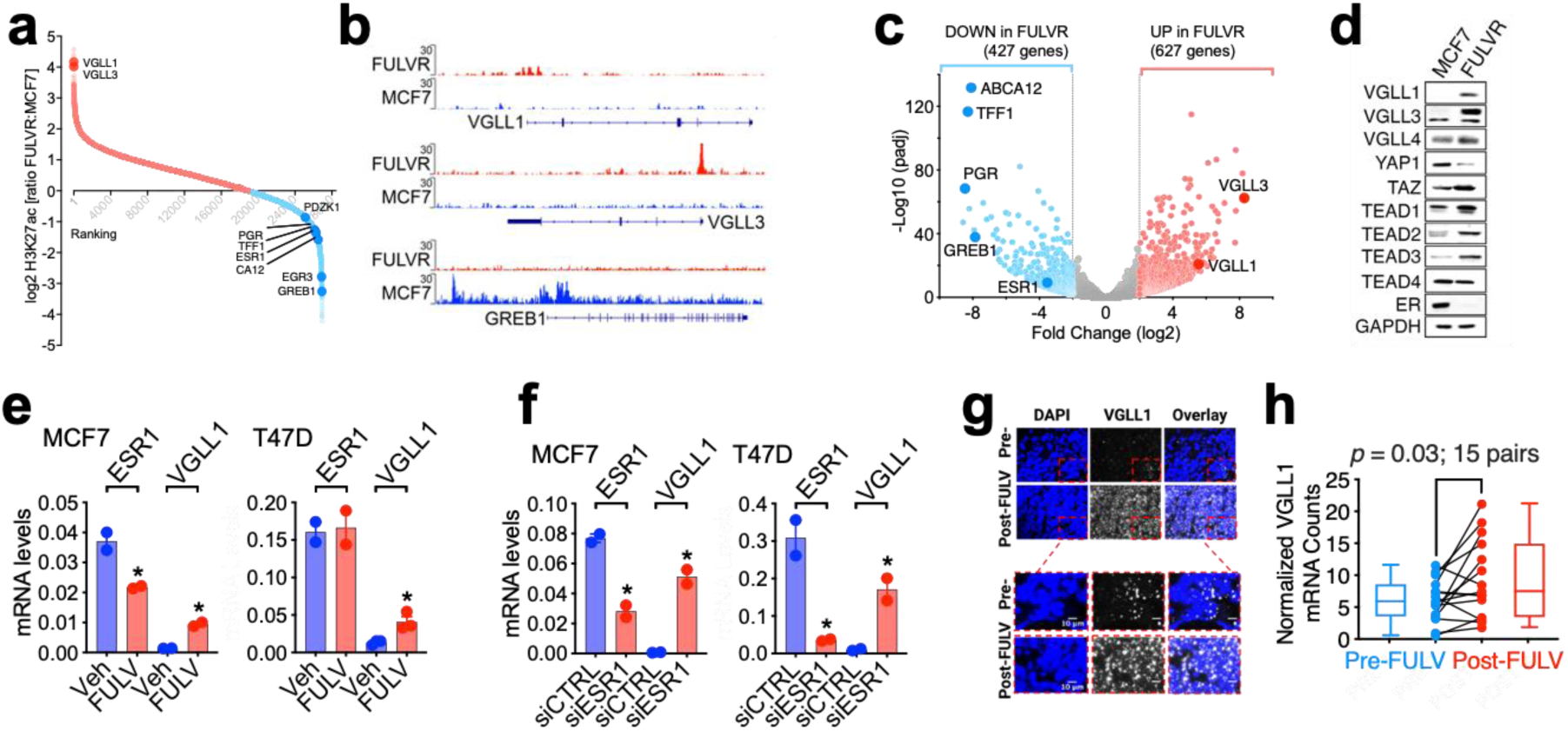
Mapping altered epigenetic landscapes in fulvestrant-resistant breast cancer identifies VGLL1. **a**, Ratio of H3K27ac in MCF7-FULVR versus MCF7 cells at gene promoters in a window of ± 1.5 kb centred on the transcriptional start site. **b**, Genome browser view of H3K27ac ChIP-seq signal at the VGLL1, VGLL3 and GREB1 genes. **c**, RNA-seq data showing upregulated genes (red, *P*adj < 0.01, log2 FC > 2) and downregulated genes (blue, *P*adj < 0.01, log2 FC < −2) in FULVR relative to MCF7 cells. **d**, Protein lysates prepared from MCF7 and FULVR cells were immunoblotted for the indicated proteins. **e**, RT-qPCR using RNA prepared from MCF7 and T47D cells following addition of 100 nM fulvestrant for 24 hours (* = p<0.05). **f,** Cells were transfected with ESR1 siRNA or a non-targeting siRNA. RNAs prepared 48 hours after transfection were used for RT-qPCR. \**P*<0.05 (Student’s t-test, two-tailed). Data are presented as mean + s.e.m. \**P*<0.05 (Student’s t-test, two-tailed). **g**-**h**, RNA-scope was performed using a probe for VGLL1 in matched pre- and post-fulvestrant treated patient samples. Representative images show the results for matched samples from one patient. Nuclei were visualised with DAPI, and individual *VGLL1* mRNA molecules detected with Cy5-labelled probes. *P*=0.03 (one-tailed Wilcoxon signed rank test). Scale bar, 10μm.

RNA-seq analysis showed that VGLL1 expression is not induced in LTED cells (Extended Data Fig. 1e), suggesting that preventing ER activation is insufficient for VGLL1 expression. By contrast, fulvestrant stimulated VGLL1 expression in ER+ breast cancer cell lines (Fig. 1e, Extended Data Fig. 1f, g). The new generation SERDs GDC0810 (Brilanestrant), AZD9496 and RAD1901 (Elacestrant) also induced VGLL1 expression (Extended Data Fig. 1h). RNAi-mediated *ESR1* knockdown promoted VGLL1 expression in all cell lines (Fig. 1f, Extended Data Fig. 1i, j), suggesting that degradation of ER protein by SERDs is important for VGLL1 induction in breast cancer cells. VGLL3, YAP, TAZ and TEAD expression was not impacted by ER knockdown. To determine if fulvestrant also promotes VGLL1 expression in tumours, we used RNAscope to measure VGLL1 mRNA in matched breast cancer biopsies taken prior to and after fulvestrant treatment. VGLL1 levels were consistently higher in post-fulvestrant samples than in the matched pre-treatment biopsies (Fig. 1g, h; Supplementary table 2). Taken together, these results clearly demonstrate that ER downregulation by SERDs induces VGLL1 in ER+ breast cancer.

### VGLL1 is recruited to TEAD binding sites at active chromatin in FULVR cells

Towards determining the role of VGLL1 in fulvestrant resistance, we performed chromatin immunoprecipitation sequencing (ChIP-seq) for identifying VGLL1 target genes. As VGLL1 is expected to be recruited via TEADs, we also profiled TEAD4, chosen since it is highly expressed in both MCF7 and FULVR cells and because successful TEAD4 ChIP-seq has been reported^24,36^. Interestingly, substantially more TEAD4 peaks were identified in FULVR cells than in the isogenic fulvestrant-sensitive MCF7 cells (20,894 vs 6,968 peaks, respectively) (Fig. 2a). Seventy-five percent of the TEAD4 binding events in MCF7 cells were also present in FULVR cells, suggesting that VGLL1 expression (and/or ER downregulation) does not stimulate redistribution of TEAD4 but rather promotes TEAD binding at many new sites. As expected, *de novo* motif analysis of the TEAD4 ChIP-seq identified TEAD binding sites as the predominant enriched motifs in both MCF7 and MCF7-FULVR cells (Fig. 2a, Extended Data Fig. 2a, b).

**Fig. 2.**
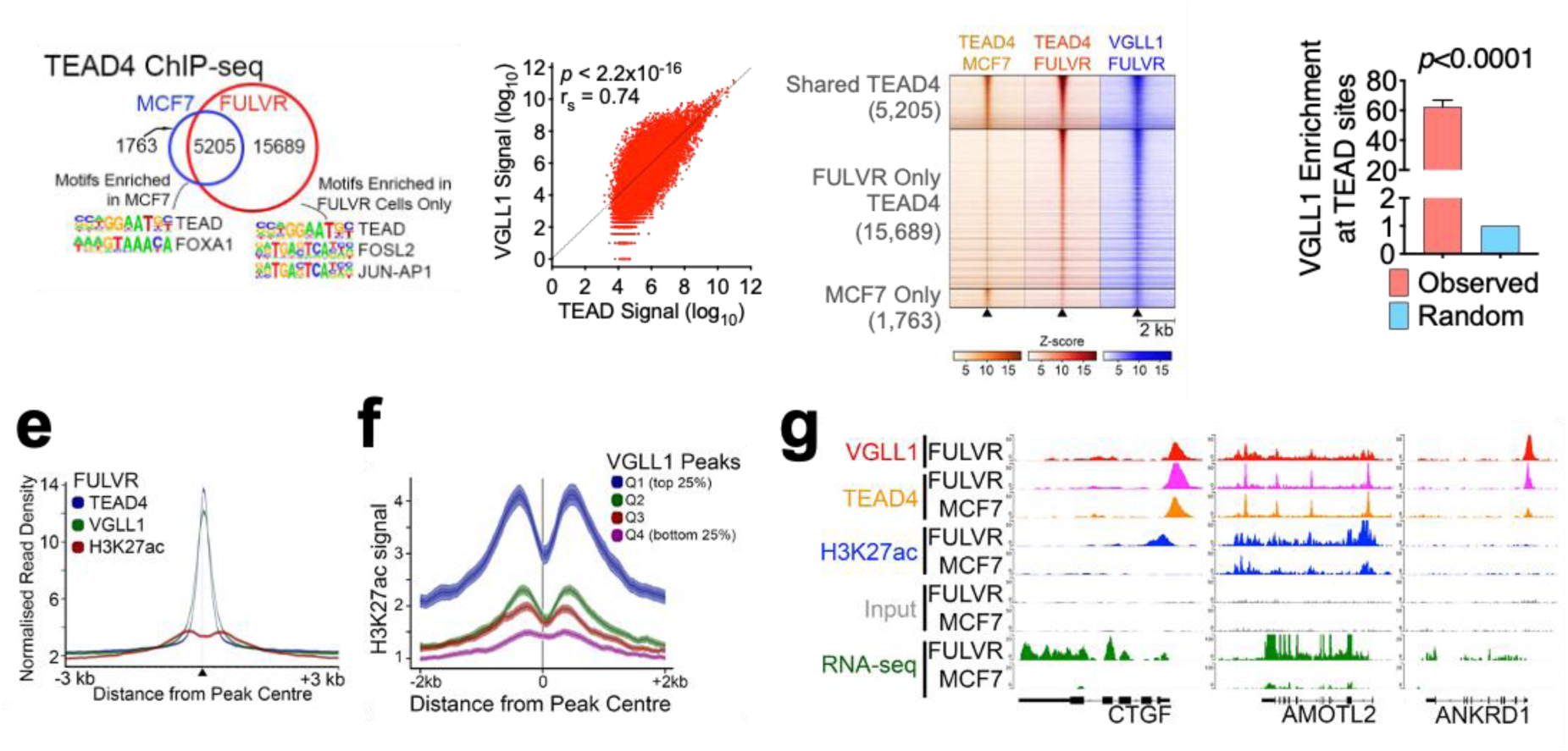
VGLL1 is recruited to TEAD4 binding regions in fulvestrant-resistant MCF7 cells. **a**, Venn diagram showing overlap between TEAD4 ChIP-seq peaks in MCF7 and FULVR cells. Top-most enriched transcription factor motifs are shown. **b**, Correlation between VGLL1 and TEAD4 occupancy at TEAD4 peaks in FULVR cells. *P*-*value* was calculated using the Spearman test; r_s_, Spearman’s correlation coefficient. **c**, Heatmap showing TEAD4 and VGLL1 binding at TEAD4 peaks common to MCF7 and FULVR (shared) and unique peaks in each cell line, in a window of ± 2kb around the peak centre. **d**, VGLL1 binding is enriched at TEAD4 peaks in FULVR cells. Binding enrichment was calculated as VGLL1-TEAD4 co-binding over the mean expected value after generating random permutations of the TEAD4 peaks (Chi-squared test p-value <0.0001). **e**, Average normalized ChIP-seq signal of TEAD4, VGLL1 and H3K27ac centred at TEAD4 peaks in FULVR cells. **f**, Average ChIP-seq signal of H3K27ac on VGLL1 peaks divided in quartiles based on the peak coverage in FULVR cells. **g**, Genome browser view of VGLL1, TEAD4 and H3K27ac ChIP-seq signal, together with the RNA-seq signal in MCF7 and FULVR cells at TEAD target genes.

Transcription activation by TEADs requires the recruitment of specific co-activators, including YAP, TAZ and VGLLs^20–22^. However, a role for VGLL1 as a TEAD co-activator has largely been restricted to reporter gene assays and co-crystallisation studies^31,32^. VGLL1 ChIP-seq of three biological replicates (Supplementary Table 3) identified 29,930 peaks common to all replicates in FULVR cells. There was a strong positive correlation between VGLL1 and TEAD4 binding events (*p*<2.2×10^-16^; Fig. 2b). VGLL1 was enriched at TEAD binding sites that were common to MCF7 and FULVR, as well as those TEAD4 binding events that were gained in FULVR cells (Fig. 2c). *De novo* motif enrichment analysis confirmed TEAD4 binding sequences as the most highly enriched sequence motif at VGLL1 binding regions (Extended Data Fig. 2c). Binding enrichment calculated as VGLL1 binding at TEAD4 peaks over random permutations of the TEAD4 binding events further confirmed VGLL1 enrichment at TEAD4 binding regions (p<0.0001; Fig. 2d).

The regions co-bound by VGLL1 and TEAD4 were present at active regulatory regions in FULVR cells, as revealed by enrichment of H3K27ac signal at regions co-occupied by VGLL1 and TEAD4 (Fig. 2e, f). Moreover, low H3K27ac signal in MCF7 cells at VGLL1/TEAD4 co-occupied regions (Extended Data Fig. 2d) is suggestive of low expression of these genes in MCF7 cells. Exemplifying this, TEAD4 was present at previously described TEAD target genes including CTGF (CCN2), AMOTL2 and ANKRD1^24^ in MCF7 and MCF7-FULVR cells (Fig. 2g), but there was substantially greater H3K27ac and expression of these genes in MCF7-FULVR cells. ChIP-qPCR confirmed that VGLL1 was recruited to these regions in MCF7-FULVR cells, but not in MCF7 cells (Extended Data Fig. 2e). YAP and TAZ were mostly absent at these genes in FULVR cells, suggesting that YAP/TAZ are not activated and that VGLL1 drives expression of TEAD-regulated genes in MCF7-FULVR cells.

### VGLL1 promotes development of fulvestrant resistance

Functional annotation of genes associated with VGLL1 binding regions showed that they are highly enriched for gene sets associated with ET resistance in breast cancer (Extended data Fig. 3a and Supplementary Table 4). Moreover, VGLL1 knockdown reduced growth of MCF7-FULVR and T47D-FULVR cells (Fig. 3a; Extended data Fig. 3b), demonstrating the functional importance of *VGLL1* in FULVR cells.

**Fig. 3.**
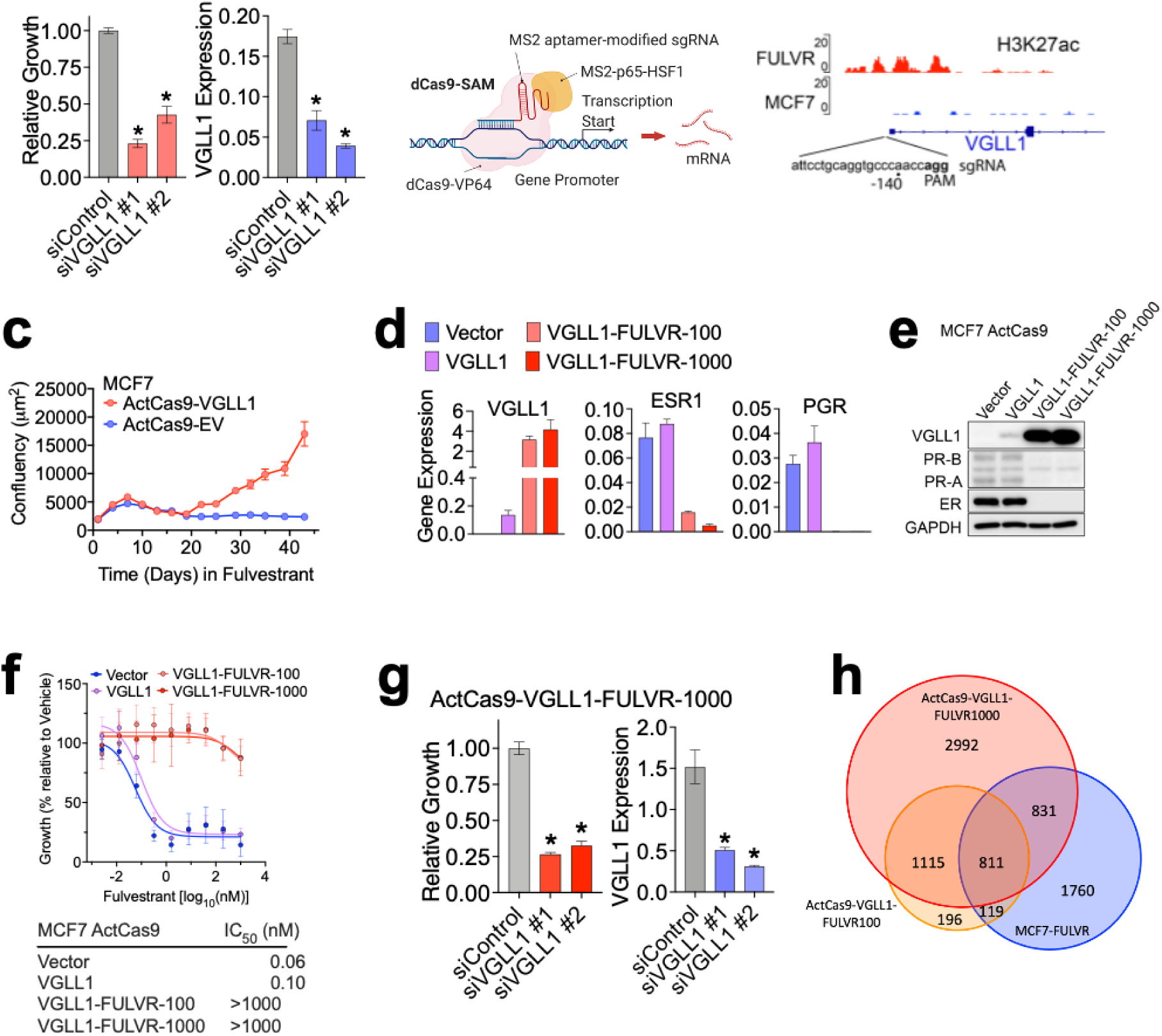
VGLL1 facilitates development of resistance to fulvestrant. **a,** MCF7-FULVR cells were transfected with two independent VGLL1 siRNAs. Growth was determined with the SRB assay five days after transfection. Data are mean ± s.e.m of n=6 independent wells. Results of one representative experiment are shown; similar results were obtained in two additional independent experiments. * *P*<0.05 (Mann-Whitney test, two tailed). Also shown are expression levels of VGLL1 following siVGLL1 transfection (*n*=3, \**p*<0.05 (Student’s t-test, two-tailed)). **b**, The synergistic activation mediator (SAM) uses a modified, catalytically dead Cas9 (dCas9), together with a sgRNA targeting to a specific gene promoter, for transcriptional activation of endogenous genes^37^. A sgRNA targeted to bp −156 to −134 of the *VGLL1* gene, was identified from the sgRNA list in ref.^37^. **c,** MCF7-ActCas9-VGLL1 cells and MCF7-ActCas9-Vector cells were cultured with 100 nM fulvestrant and cell confluency was measured using Incucyte live cell imaging. Data show mean ± sem of n=9 representative images. **d,** RNA prepared from the indicated cells was used for RT-qPCR (mean ± sem; n=3). **e,** Immunoblotting of cell lysates prepared from the indicated cell lines. VGLL1-FULVR-100 and VGLL1-FULVR-1000 are fulvestrant resistant cell lines derived from the parental MCF7 ActCas9-VGLL1 cell line after continuous culturing in the presence of either 100 nM or 1000 nM fulvestrant, respectively. **f,** Growth of the indicated cell lines treated with increasing concentrations of fulvestrant to a maximum of 1 µM, for five days. Cell growth was estimated using the SRB assay and is shown as percentage relative to vehicle (n=6). Half maximum inhibitory concentration (IC_50_) is indicated. **g,** MCF7 ActCas9-VGLL1-FULVR cells were transfected with siVGLL1 and growth assessed as in **a.** Data are mean ± s.e.m of n=6. One representative experiment is shown; similar results were obtained in two additional independent experiments. * *P*<0.05 (Mann-Whitney test, two tailed). RT-qPCR for VGLL1 is also shown (*n*=3, \**p*<0.05 (Student’s t-test, two-tailed)). **h,** Venn diagram comparing differentially expressed genes in MCF7-FULVR and MCF7-ActCas9-VGLL1-FULVR cells. 5,749 genes differentially expressed (padj<0.05) between MCF-FULVR and fulvestrant-sensitive MCF7 cells. 2,241 differential genes were identified in MCF7-ActCas9-VGLL1-FULVR-1 versus MCF7-ActCas9-VGLL1. There were 6,923 differential genes in MCF7-ActCas9-VGLL1-FULVR-2 relative to MCF7-ActCas9-VGLL1 cells.

To determine if *VGLL1* is sufficient for the development of resistance to fulvestrant, we induced expression of the endogenous *VGLL1* gene using the CRISPR/Cas9 Synergistic Activation Mediator (SAM)^37^ targeted to the *VGLL1* gene promoter (Fig. 3b). Although VGLL1 expression was induced (ActCas9-VGLL1; Extended data Fig. 3c-d), this, in of itself was insufficient for an altered response to fulvestrant (see below). Treatment of ER+ breast cancer cells with fulvestrant causes rapid growth arrest and the establishment of resistance requires prolonged culturing with fulvestrant over many months^38–40^. In agreement with these reports, growth inhibition was maintained in vector control cells (MCF7-ActCas9-Vector) for >100 days in the presence of 100 nM fulvestrant (Fig. 3c, Extended data Fig. 3e). By contrast, following an initial period of stasis, growth recovery was observed in fulvestrant-treated cells within just 20 days in VGLL1 expressing cells. Interestingly, development of fulvestrant resistance in these cells was accompanied by further increases in levels of VGLL1, together with reduced ER and PR expression (Fig 3d-e). Similar results were obtained for ActCas9-VGLL1 cells following treatment over 100 days with 1000 nM fulvestrant (MCF7 Act-Cas9-VGLL1-FULVR1000) cells. MCF7-ActCas9-VGLL1-FULVR cells were cross-resistant to next generation SERDs (Fig. 3f, Extended data Fig. 3f). Dependence of the ActCas9-VGLL1-FULVR cells on VGLL1 was confirmed with VGLL1 siRNA (Fig. 3g).

Interestingly, induction of VGLL1 (MCF7-ActCas9-VGLL1) was not accompanied by marked changes in gene expression (Extended data Fig. 3g). By contrast, 2,241 and 3,521 genes were substantially altered in the two fulvestrant-resistant ActCas9-VGLL1 cell lines (Fig. 3h). The majority (85%) of differential genes in ActCas9-VGLL1-FULVR100 cells were also significantly altered in ActCas9-VGLL1-FULVR1000 cells. Moreover, of the 360 genes up-regulated with log2FC≥1 in MCF7-FULVR cells, most were also up-regulated (log2FC≥1) in ActCas9-VGLL1-FULVR100 (341/360 (94%) genes) and ActCas9-VGLL1-FULVR1000 cells (338/360 (94%) genes) (Supplementary Table 5, Extended data Fig. 3h). A similar relationship was observed for the 175 downregulated genes (log2FC≤-1) in MCF7-FULVR, wherein 90% of these genes were also downregulated in the ActCas9-VGLL1-FULVR cells.

Our results show that although expression of VGLL1 alone is insufficient for resistance to fulvestrant, its expression promotes progression to fulvestrant resistance. It is possible that VGLL1 levels above a threshold achieved with ActCas9 targeting are needed for resistance to fulvestrant. Indeed, VGLL1 levels in the resistant cells were considerably higher than those in the MCF7-Act-Cas9-VGLL1 cells. Alternatively, other genes induced or repressed by prolonged fulvestrant treatment may be necessary for VGLL1 activation. One such requirement might be TEAD1, given that its expression is commonly elevated in the different fulvestrant-resistance models (Supplementary Table 5). Another possibility is that post-translational modification of VGLL1 controls its activity. Indeed, the levels and activities of the other major TEAD partners, YAP and TAZ, are critically dependent on their phosphorylation by the HIPPO pathway LATS1/2 kinases^41^. However, the lack of a LATS recognition motif in VGLL1 ^31^ makes it unlikely that VGLL1 is regulated by the HIPPO pathway. Taken together, downregulation of ER and its transcriptional programmes by fulvestrant appears to be a necessary first step for adaptation of ER+ breast cancer cells away from ER dependence to a requirement for VGLL1.

### Transcriptional dependency of FULVR cells on VGLL1

For understanding the importance of VGLL1 in directing gene expression in fulvestrant resistant cells, we first stratified genes based on the presence or absence of VGLL1 ChIP-seq peaks. Expression of genes that are bound by VGLL1 (n=3,195) was significantly higher in FULVR than in fulvestrant-sensitive MCF7 cells (Fig. 4a). The genes associated with VGLL1 peaks were also more highly expressed in ActCas9-VGLL1-FULVR cells (Extended data Fig. 4a). To further assess VGLL1-regulated gene expression in MCF7-FULVR cells, we performed RNA-seq following VGLL1 knockdown using 3 VGLL1 siRNAs. Robust VGLL1 knockdown with each VGLL1 siRNA was confirmed in each biological replicate RNA (n=3; Extended data Fig. 4b). Expression of 3500-4000 genes was significantly (padj < 0.05) altered by each VGLL1 siRNA (Extended data Fig 4c), with around half of these genes being downregulated. To identify direct VGLL1 targets we filtered genes that were downregulated with at least 2 independent siRNAs (1,372 genes, padj < 0.05) for the presence of VGLL1 ChIP-seq peaks within 2.5 kb of transcription start sites (TSS). This defined 762 genes, which we termed VGLL1-activated genes. Not-VGLL1 targets, so-called because they were unaffected by VGLL1 knockdown and did not contain VGLL1 peaks, number 8,932. Comparing the VGLL1-activated genes showed that they were also more highly expressed in MCF7-FULVR cells than in the isogenic fulvestrant-sensitive cells, whereas there was no difference in levels of the not-VGLL1 targets (Fig. 4b). As expected, VGLL1-regulated genes were also enriched for TEAD4 binding regions (Extended data Fig. 4d, e) and were associated with low H3K27ac at TSS in MCF7 cells but high H3K27ac in MCF7-FULVR cells (Fig 4c, Extended data Fig. 4f, g). Functional annotation of the VGLL1-activated genes revealed enrichment for pathways associated with growth factor signalling, including growth factor binding and transmembrane receptor protein kinase activity (Fig. 4e), exemplified by IGFBP3, ITGB6, TGFB2 and EGFR (Fig. 4f, g; Extended data Fig. 4h). Taken together, our findings strongly support the premise that VGLL1/TEAD is a key driver of genes whose expression is increased in progression of ER+ breast cancer cells to fulvestrant resistance.

**Fig. 4.**
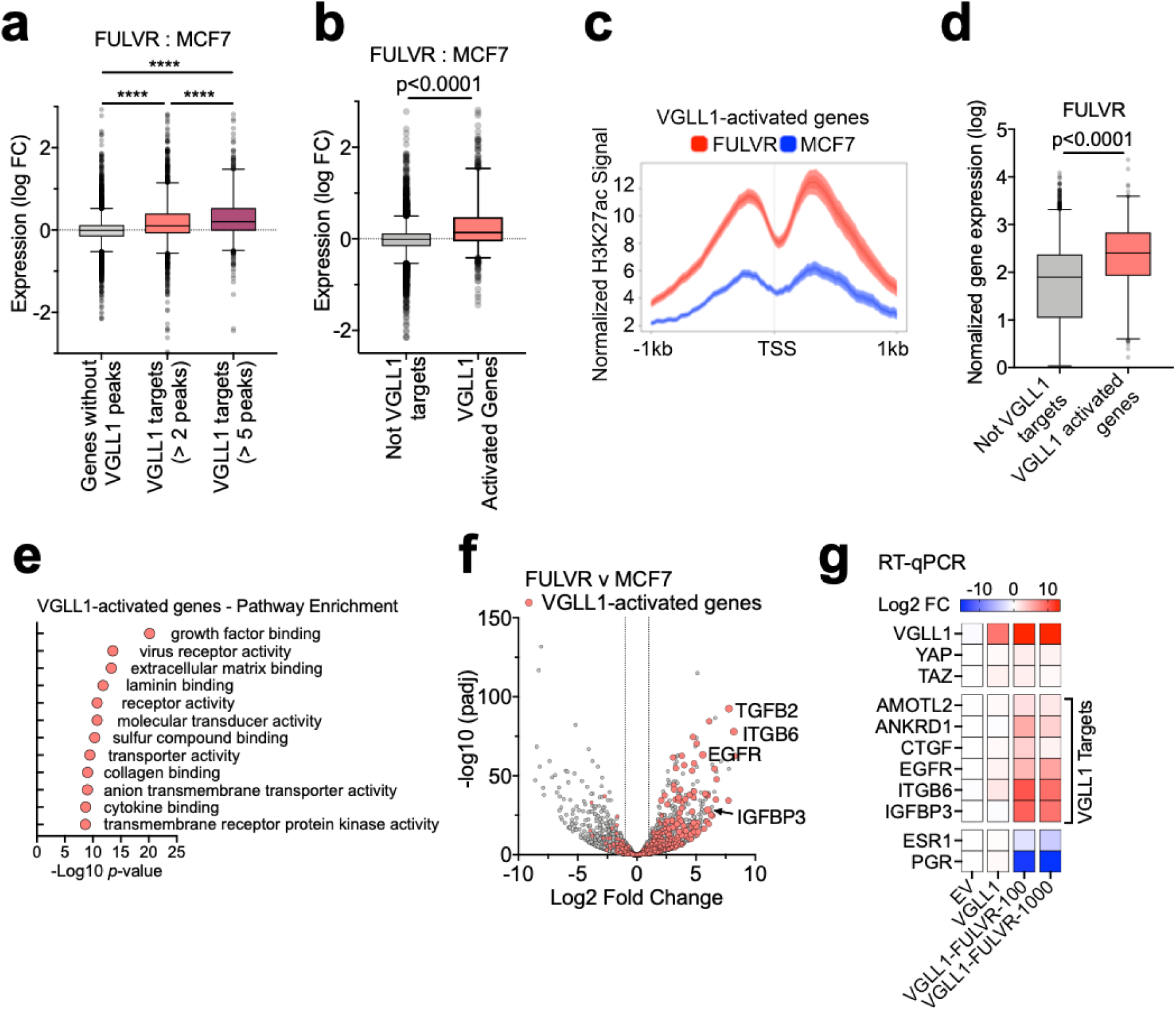
Genes upregulated in fulvestrant-resistant breast cancer cells depend on VGLL1 transcriptional activity. **a,** Genes predicted as VGLL1 targets in FULVR cells are more highly expressed in FULVR cells than in MCF7 cells. Genes were segregated into those with no VGLL1 peaks, and those with 2-4 and ≥5 VGLL1 peaks. The y-axis shows the log2 fold change in gene expression determined from RNA-seq in FULVR cells versus the parental MCF7 cells. \*\*\*\**P*<0.0001 (Mann-Whitney test, two tailed). **b,** Genes activated by VGLL1 (*n* = 762) are over-expressed in FULVR cells relative to MCF7 cells, compared to not-VGLL1 targets (*n* = 8,932). VGLL1-activated genes and not-VGLL1 targets were determined by RNA-seq in FULVR cells transfected with VGLL1 siRNAs. The VGLL1-activated genes were defined as genes downregulated by VGLL1 siRNAs (*P*<0.0001 (Mann-Whitney test, two tailed)). **c,** Normalized average H3K27ac signal on the promoters of VGLL1 activated genes in FULVR cells and MCF7 cells. **d,** VGLL1-activated genes (*n* = 762) are more highly expressed in FULVR cells than the not-VGLL1 targets (*n* = 8,932). The y-axis shows normalized gene expression values from RNA-seq. *P*<0.0001 (Mann-Whitney test, two tailed). **e,** GO molecular function sets enriched in VGLL1-activated genes. **f,** RNA-seq data represented as a volcano plot for the comparison between FULVR vs MCF7 cells. **g,** RT-qPCR was performed using RNA prepared from the indicated cell lines. Gene expression was normalized to *GAPDH* expression and is shown as log2 fold difference relative to expression in MCF7 ActCas9-Vector cells.

### VGLL1 induces EGFR expression in breast cancer cells to drive growth of FULVR cells

The above findings link high VGLL1 activity with genes implicated in receptor tyrosine kinase (RTK) signalling, including EGFR. High expression of EGFR and/or its downstream effectors has been linked to reduced efficacy of ET, including tamoxifen and fulvestrant^42^. Furthermore, ectopic expression of EGFR demonstrably promotes fulvestrant resistance in breast cancer cells^1^. Indeed, EGFR expression was elevated in all our FULVR cell lines compared with their isogenic fulvestrant-sensitive counterparts (Extended data Fig. 5a). Analysis of our ChIP-seq data revealed gain of TEAD4 binding at the EGFR gene in MCF7-FULVR cells, particularly at a region ∼50 kb upstream of the transcription start site where it was co-localised with VGLL1 (Fig. 5a). Enhanced TEAD4 binding at this region was confirmed by ChIP-qPCR in MCF7-ActCas9-VGLL1-FULVR cells, was accompanied by the presence of VGLL1 in this region and showed induction of H3K27 acetylation (Fig. 5b). VGLL1 and TEAD4 co-binding at this region was consistent with the substantial increases in EGFR levels in MCF7-ActCas9-VGLL1-FULVR cells (Fig. 5c). Immunoblotting confirmed EGFR over-expression in MCF7-ActCas9-VGLL1-FULVR cells, accompanied by greater activation of downstream effectors AKT and ERK1/2 MAPK (Fig. 5d).

**Fig. 5.**
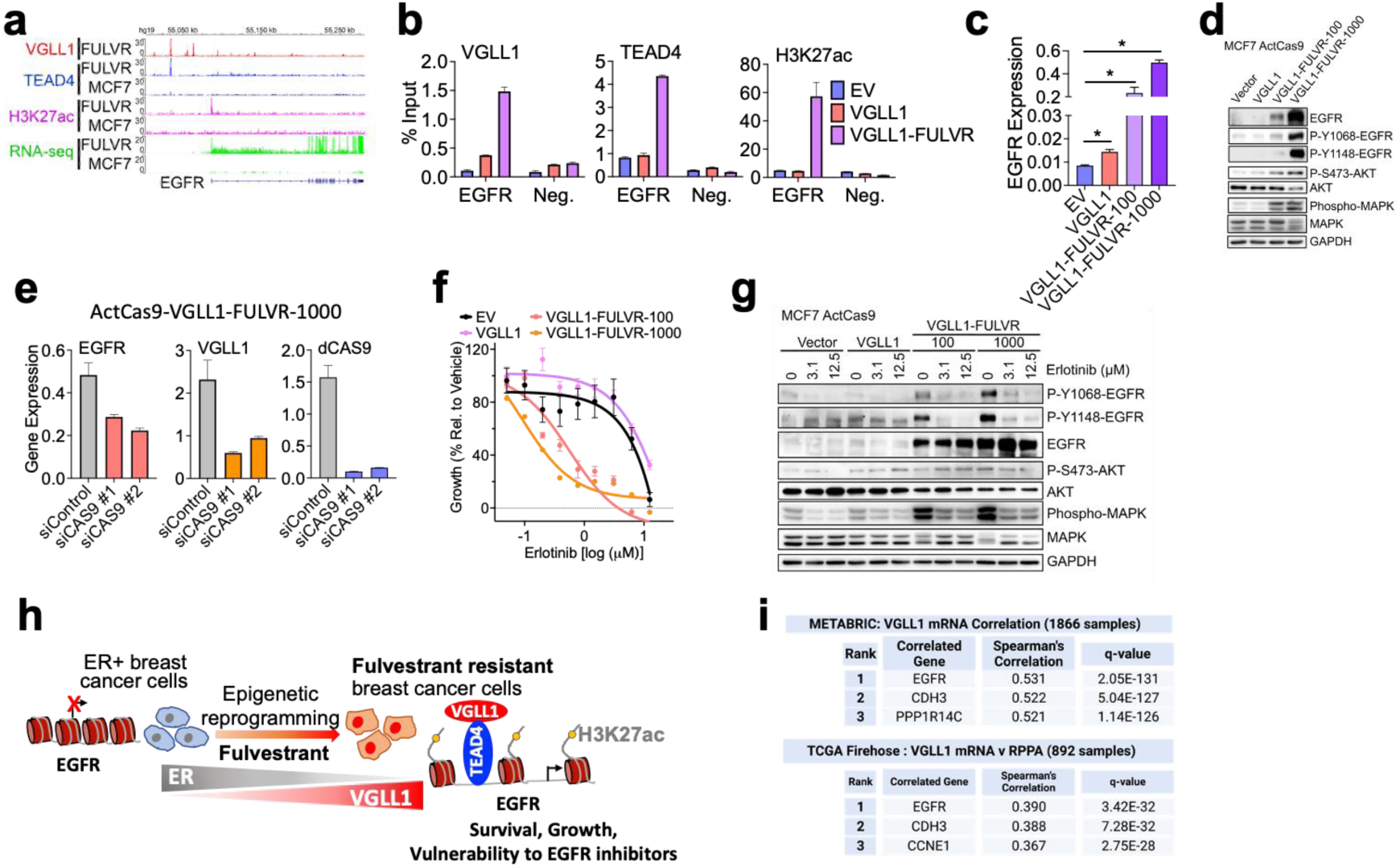
VGLL1 induces EGFR expression and sensitivity to EGFR inhibition in FULVR cells. **a,** Genome browser view of VGLL1 and TEAD4 ChIP-seq signal at the EGFR gene and EGFR enhancer (highlighted). **b,** ChIP-qPCR for TEAD4, VGLL1 and H3K27ac in MCF7 ActCas9-Vector, MCF7 ActCas9-VGLL1 and MCF7 ActCas9-VGLL1-FULVR cells showing VGLL1 and TEAD4 binding at the EGFR enhancer together with EGFR enhancer activation exclusively in FULVR cells. The CTGF −8.3kb region was used as a negative control for VGLL1/TEAD4 binding. **c,** RT-qPCR for EGFR in the indicated MCF7-ActCas9 cells (n=3, *p<0.05 (Student’s t-test, two-tailed)). **d,** Immunoblotting for EGFR and downstream EGFR signalling proteins showing EGFR up-regulation and activation of the EGFR pathway in MCF7 ActCas9-VGLL1-FULVR cells. **e,** RT-qPCR for dCAS9, VGLL1 and EGFR in MCF7 ActCas9-VGLL1-FULVR cells transfected with two independent dCAS9 siRNAs show reduction in VGLL1 and EGFR expression. **f,** Growth of the indicated MCF7-ActCas9 cells treated with increasing concentrations of the EGFR inhibitor erlotinib (0.05 μM to 12.5 μM) for 5 days. Growth is shown as percentage relative to vehicle treatment. **g,** Western blot of EGFR and the downstream EGFR signalling proteins in the indicated MCF7 ActCas9 cell lines treated with erlotinib at the indicated concentrations (24 h). **h**, Model for a mechanism by which inhibition of ER activity and concomitant VGLL1 induction drive EGFR expression in fulvestrant-resistant breast cancer cells, to promote cell survival and growth. **i,** EGFR ranks as the most significantly co-expressed gene with VGLL1 in breast cancer patients from METABRIC and is also the highest ranked protein correlated with VGLL1 in breast cancer from TCGA Firehose legacy cohort. In each case the top 3 highest ranked genes are shown. The ranking and correlations were generated from cBioportal (accessed 15/12/2022).

Since VGLL1 expression in MCF7-ActCas9-VGLL1-FULVR cells was induced by promoter targeting of ActCas9, we transfected these cells with siRNAs for Cas9, which reduced VGLL1 expression and lowered EGFR expression (Fig. 5e). Similarly, VGLL1 knockdown was sufficient to reduce levels of EGFR in the FULVR cells generated by long-term culturing with fulvestrant (Extended data Fig. 5b-d). Consistent with high EGFR expression and activity, treatment of MCF7-ActCas9-VGLL1-FULVR cells with the EGFR inhibitor erlotinib^43^, inhibited cell growth, accompanied by reductions in AKT and MAPK phosphorylation (Fig. 5f, g). By contrast, the isogenic fulvestrant-sensitive cells were substantially less sensitive to erlotinib. A similar difference in sensitivity to erlotinib was observed for MCF7-FULVR and ZR-75-1-FULVR cells compared with the isogenic fulvestrant-sensitive MCF7 cells (Extended data Fig. 5e-f).

Our results reveal a mechanism explaining the induction of EGFR in ET-resistant breast cancer, in which epigenomic remodelling due to ER downregulation results in the induction of VGLL1 and consequent VGLL1-TEAD4 co-binding at the *EGFR* enhancer to induce EGFR expression (Fig. 5h). In support of our findings, EGFR expression was positively correlated with VGLL1 expression in ER+ breast cancer patients in the TCGA^44^ and METABRIC^45^ cohorts (Extended data Fig. 5h). Remarkably, EGFR was the highest-ranked gene co-expressed with VGLL1 at the mRNA level in METABRIC, as well as the highest ranked protein significantly correlated with VGLL1 in breast cancer in the TCGA series (Fig. 5i). These analyses together corroborate our cell line studies in identifying a direct role for VGLL1 in driving EGFR expression in breast cancer.

### Pharmacological inhibition of VGLL1-TEAD interaction in fulvestrant-resistant breast cancer

The small molecule drug verteporfin (VP) inhibits TEAD-dependent gene expression by blocking its interaction with YAP, resulting in downregulation of YAP/TEAD target genes^46,47^ Given the structural similarities between YAP-TEAD and VGLL1-TEAD complexes^31,32^, we reasoned that VP would also disrupt VGLL1-TEAD4 interaction and so inhibit expression of VGLL1-regulated genes. Consistent with this model, treating FULVR cells with VP inhibited their growth (Fig. 6a, Extended data Fig. 6a). VP reduced recruitment of VGLL1 to target genes including EGFR, whereas TEAD4 binding was unaffected or only modestly reduced by VP (Fig. 6b), demonstrating that VP prevents recruitment of VGLL1 to TEAD4 binding regions.

**Fig. 6.**
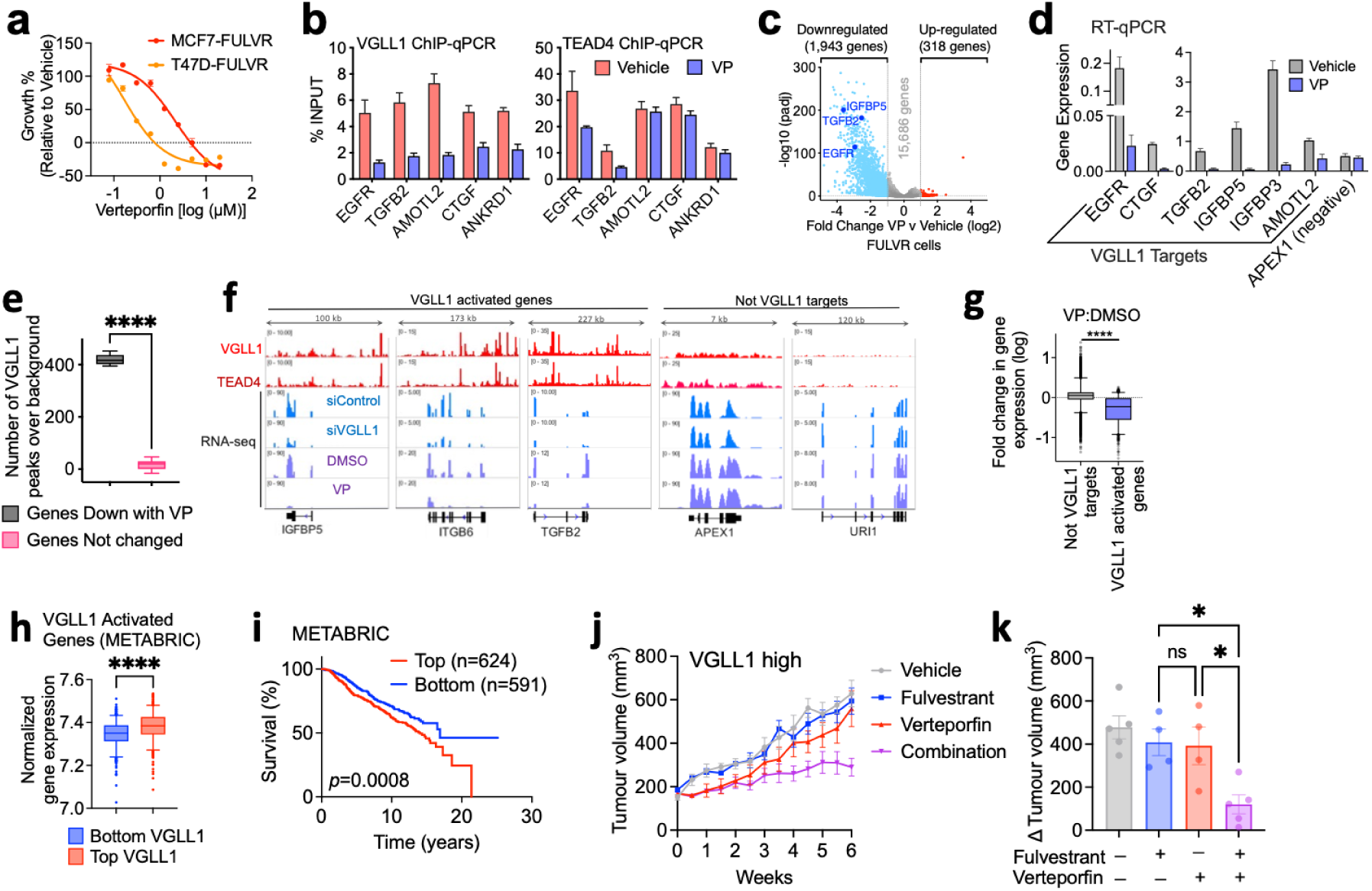
Verteporfin inhibits VGLL1 transcriptional activity and resensitises breast cancer cells to fulvestrant. **a,** VP impairs the growth of FULVR cells. Growth in the indicated FULVR cells treated with increasing concentrations of VP is shown as percentage of growth relative to vehicle. **b,** ChIP-qPCR for VGLL1 and TEAD4 in MCF7-FULVR cells showing reduced VGLL1 binding at the target genes in the presence of VP (2μM, 24 h). The y-axis shows DNA enrichment calculated as the percentage of input. **c,** Volcano plot of RNA-seq data (from *n*=4 biological replicates) showing upregulated genes (red, adjusted *P*<0.01, log2 (fold change) > 1) and downregulated genes (blue, adjusted *P*<0.01, log2 (fold change) < −1) in MCF7-FULVR cells treated with VP, (2μM, 24 h) compared to vehicle (DMSO). **d,** RT-qPCR in MCF7-FULVR cells treated with VP (2μM, 24 h) showing that VP selectively downregulates the expression VGLL1 targets, while the expression of the not-VGLL1 target (APEX1) is not affected by VP. **e,** Genes downregulated by VP are highly enriched in VGLL1 peaks. The y-axis shows the number of VGLL1 peaks over the expected value after generating random permutations of the VGLL1 peaks showing that the top-most downregulated genes after VP treatment in FULVR cells (*n* = 631) have significantly higher number of VGLL1 peaks compared to the bottom genes not differentially expressed by VP (*n* = 631). Data are presented as box-and-whiskers plots (whiskers extend from the 5th to the 95th percentile; the box extends from the 25th to the 75th percentile; the line within the box represents the median). **** *P*<0.0001 (Mann-Whitney test, two tailed). **f,** Genome browser view of VGLL1 and TEAD4 normalized ChIP-seq signal and normalized RNA-seq signal in FULVR cells showing representative examples of VGLL1 activated genes (direct VGLL1 targets) and not-VGLL1 targets. **g,** VGLL1 activated genes (*n* = 762) are preferentially downregulated by VP compared to not-VGLL1 targets (*n* = 8,932). The y-axis shows fold change in gene expression from RNA-seq between VP versus DMSO treatment in MCF7-FULVR cells. \*\*\*\**P*<0.0001 (Mann-Whitney test, two tailed). **h,** Breast cancer patients with higher *VGLL1* expression display increased levels of expression of the VGLL1 activated genes compared to patients with lower VGLL1 expression. Patients from the METABRIC breast cancer dataset were stratified according to high (top quantile, n=495) or low (bottom quantile, n=495) *VGLL1* expression levels. Data are presented as in **e**. \*\*\*\**P*<0.0001 (Mann-Whitney test, two tailed). **i,** Kaplan–Meier plot representing the percentage of metastasis-free survival in patients with ER+ breast cancer patients treated with endocrine therapies showing that patients with higher expression of the VGLL1 activated genes signature display lower survival rates (log-rank Mantel-Cox test). **j,** Tumour growth curves for mice bearing the KCC_P_3837 FulvR clone 2 (VGLL1 high) PDX model, treated as shown. **k,** Mean tumour volumes at 6 weeks; dots show sizes of the individual tumours. * = two-tailed unpaired T test, P < 0.05.

For global assessment of the inhibition of VGLL1 co-transcriptional activity by VP we performed RNA-seq in MCF7-FULVR cells following the addition of VP. VP potently downregulated the expression of VGLL1 activated genes, including EGFR (Fig. 6c), findings that could be confirmed with RT-qPCR (Fig. 6d). These results are consistent with reduced expression of these genes following VGLL1 knockdown (Extended data Fig. 6b, c; see also Fig. 5). Genes downregulated by VP were enriched for VGLL1 peaks (Fig. 6e). Indeed, expression of genes that contained promoter proximal VGLL1 peaks was significantly reduced by VP, compared to lack of change in expression for genes without VGLL1 binding (Extended data Fig. 6d). The selectivity of VP in downregulating VGLL1 target genes was further confirmed by analysing all the high confidence VGLL1 targets and not-VGLL1 targets, revealing that VGLL1 activated genes were significantly downregulated by VP, compared to not-VGLL1 targets (Fig. 6f, g). Indeed, the majority of VGLL1 activated genes were downregulated by VP and displayed preferential sensitivity to VP (Extended data Fig. 6e). In line with this, we also found that genes that were downregulated by VP were expressed at higher levels in FULVR cells, compared to genes not inhibited by VP (Extended data Fig. 6f), consistent with VGLL1 activated genes being more highly expressed than not-VGLL1 targets in FULVR cells (Fig. 4d). Finally, genes downregulated by VP were enriched in functional categories attributed to VGLL1 activated genes including growth factor binding, extracellular matrix binding and transmembrane receptor protein kinase activity (Extended data Fig. 6g and Supplementary Table 6).

Interestingly, expression of the 762 genes identified as likely VGLL1 activated genes was positively associated with VGLL1 levels in the METABRIC breast cancer cohort (Fig 6h). Moreover, when we stratified patients according to the combined expression levels of the VGLL1 activated genes, we found that ER+ patients with high expression levels of this gene set had worse prognosis (Fig. 6i), indicating that our signature of VGLL1 activated genes may have prognostic value for breast cancer patients treated with endocrine therapies. Overall, our results show that VGLL1 transcriptional activity is sensitive to VP treatment and suggest that inhibition of VGLL1 recruitment to TEADs could provide an approach to treating VGLL1-dependent, endocrine-resistant breast cancer.

To test the efficacy of VP *in vivo*, we used a patient-derived xenograft (PDX) model, KCC_P_3837, generated from an untreated grade 3, ER-positive, PR-positive, HER2-negative primary invasive ductal carcinoma, in which fulvestrant-resistant derivatives were generated through serial passaging in mice. To directly assess the role of VGLL1 *in vivo*, we treated one fulvestrant-resistant clone having high VGLL1 expression and a second that had low/absent VGLL1 expression, with VP alone or in combination with fulvestrant (Extended data Fig. 6i). Interestingly, while VP treatment did not impact growth of the VGLL1-low PDX, combination of VP with fulvestrant significantly reduced growth of the VGLL1-high PDX (Fig. 6j-k, Extended data Fig 6j-k).

## Discussion

Our study has revealed that VGLL1 expression is induced by endocrine therapies that cause ER down-regulation. We further show that ER down-regulation by these drugs drives the epigenetic reprogramming that permits the establishment of VGLL1-directed transcriptional programs that support ER-independent cell survival and proliferation. These VGLL1-directed transcriptional programs involve its recruitment to TEAD binding regions. Indeed, reprograming of the epigenetic landscape in breast cancer cells resistant to fulvestrant is associated with increased TEAD4 recruitment to the chromatin and association of VGLL1 and TEAD4 co-binding with active enhancers in the resistant cells.

The well-established oncogenic role of TEADs in different cancers has been closely linked to the activities of YAP and TAZ as TEAD coactivators^48,49^. However, it is increasingly clear that the canonical functions of TEADs as downstream effectors of YAP and TAZ in the Hippo pathway do not provide the sole mechanism by which TEADs contribute to cancer progression in different cancers^36,50,51^. For instance, in ER+ breast cancer, a non-canonical function of YAP and TEAD4, has been reported, wherein they act as ER cofactors, regulating ER target genes rather than canonical TEAD target genes^36^. Our results agree with these findings, as we found that canonical TEAD target genes were expressed only at low levels in ER+ breast cancer cells and we did not observe enrichment of YAP/TAZ binding at these genes. By contrast, many of the established TEAD target genes^52–54^ are upregulated in fulvestrant-resistant cells, yet YAP/TAZ binding is largely absent at these genes. Instead, VGLL1 was co-bound with TEAD at these genes in FULVR cells and the expression of these genes was dependent on VGLL1.

Demonstration of the importance of VGLL1/TEAD in promoting resistance to ER down-regulators also permits identification of targetable pathways engaged by VGLL1/TEAD, as exemplified herein for EGFR. EGFR expression was greatly elevated in FULVR cells, accompanied by enhanced TEAD4 binding proximal to the EGFR gene. VGLL1 was co-bound with TEAD4 at this region and EGFR expression in fulvestrant-resistant cells was dependent on VGLL1. ERK1/2 MAPK and AKT activities were elevated in FULVR cells, these activities being dependent on EGFR activity as demonstrated by the inhibition observed with erlotinib. Consistent with these results, FULVR cells were substantially more strongly growth inhibited by erlotinib than the fulvestrant-sensitive cells. VGLL1 knockdown reduced EGFR, as well as lowering phospho-AKT levels, indicative of the importance of VGLL1 for EGFR expression and activity in FULVR cells. Supporting our findings is the observation that EGFR is the most highly ranked gene co-expressed with VGLL1 in breast cancer samples. Recent studies show that EGFR and HER2 amplification, as well as mutations of genes in the downstream MAPK pathway are increased in metastatic, endocrine-resistant ER+ breast cancer, together accounting for 10-15% of cases^1,55^ and demonstrating the importance of elevated EGFR signalling in acquired resistance to endocrine treatments. Our results evidence an alternate mechanism by which EGFR expression can be induced through VGLL1 expression resulting from endocrine therapy, leading to endocrine-resistant breast cancer and thus advance a rationale for an expanded clinical utility for EGFR inhibitors in advanced ER+ breast cancer.

While direct inhibition of VGLL1 would provide the most effective approach for targeting VGLL1-expressing breast cancers, no VGLL1 inhibitors are currently available, whereas several small molecule TEAD inhibitors have been reported^56–58^. One of these, verteporfin, an FDA approved photodynamic therapy drug for macular degeneration, which serendipitously disrupts YAP/TEAD interactions^46,47^, prevented VGLL1 recruitment to TEAD binding sites and inhibited expression of VGLL1 target genes. Verteporfin re-sensitised a VGLL1-expressing fulvestrant-resistant PDX model to fulvestrant *in vivo*. Thus, our findings show that VGLL1 transcriptional and growth dependencies could be exploited as a therapeutic vulnerability in advanced ER+ breast cancer and so inhibiting VGLL1 interaction with TEAD or inhibition of downstream VGLL1-activated genes such as EGFR, could be viable therapeutic options for patients who progress on ER-downregulating endocrine therapies.

## Acknowledgements

This work was funded through Cancer Research UK grants C37/A9335, C37/ A12011, and C37/A18784. Additional support was provided by the Imperial College Healthcare NHS Trust Tissue Bank, the Imperial Experimental Cancer Medicine Centre and the Imperial NIHR Biomedical Research Centre. The views expressed are those of the authors and not necessarily those of the NHS, the NIHR or the Department of Health.

## Author contributions

C.G., L.B, R.C.C and S.A. designed the experimental work. C.G., C-F. L, A.K.S, M.P., A.J.N, X.W. and G.M.S. undertook the experimental work. A.B. and G.P. provided the patient samples. C.M.D, N.P. S. Abuelmaaty and E.R. assessed the patient samples. C.G., H.F and V.T.M.N undertook the bioinformatics analyses. V.M.T.N. and L.M provided the ChIP-seq and RNA-seq data for the fulvestrant-resistant cells. L-A.M. provided additional fulvestrant-resistance cell line models. The *in vivo* study was designed and undertaken by N.P., H.Z.M and E.L. C.G and S.A. drafted the first version of the manuscript, and all authors contributed to its final form.

---

**Extended Data Figure 1.**
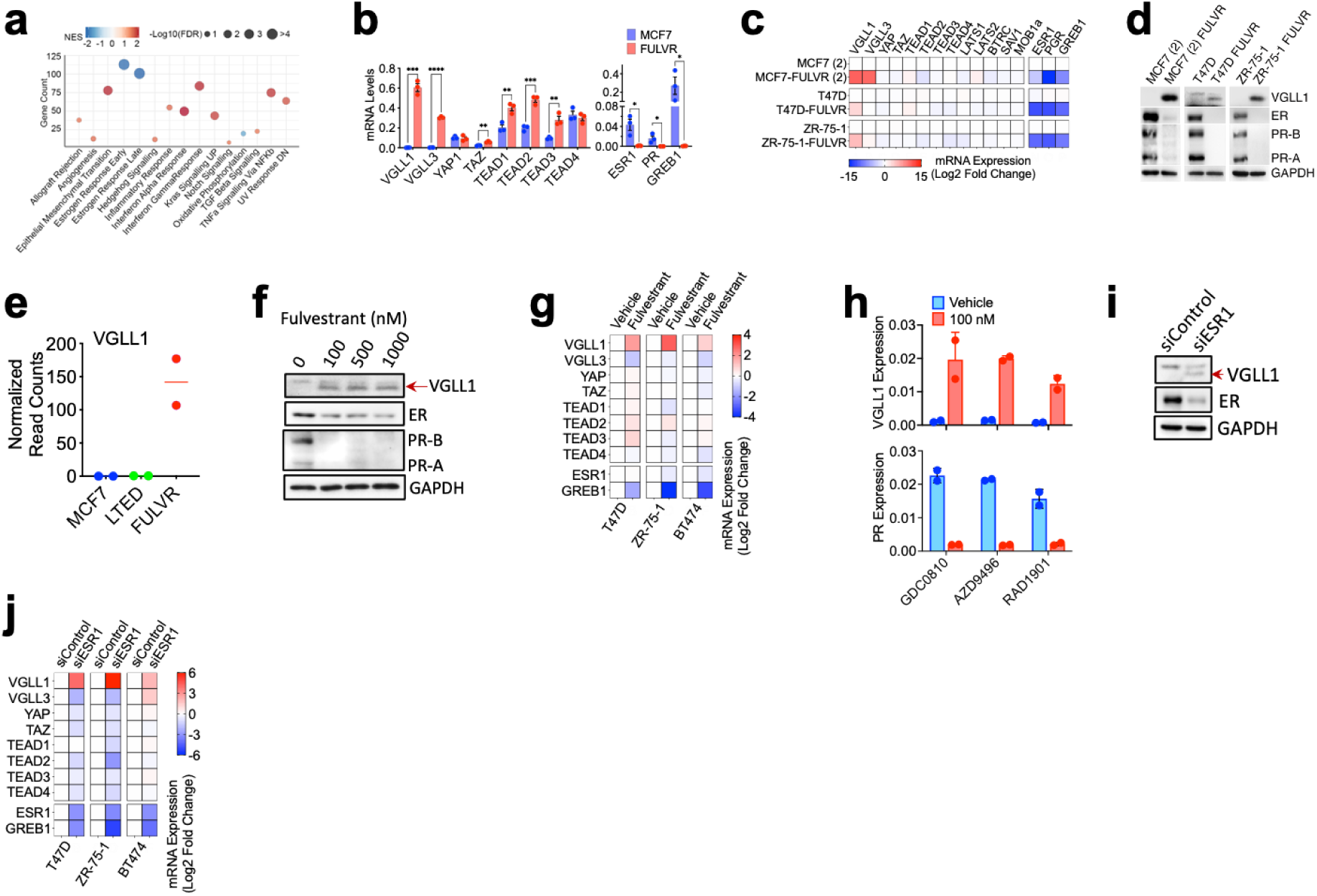
VGLL1 expression is induced by SERD-mediated ER downregulation. **a,** Gene set enrichment analysis of RNA-seq data for the MSigDB hallmark gene sets significantly enriched in MCF7-FULVR cells relative to the isogenic MCF7 cells. The bubble plot was generated using the R package ‘gplot’. **b,** RT-qPCR for genes in the YAP/TEAD pathway in MCF7 and FULVR cells. Data are presented as mean, error bars = s.e.m, from n=3 experiments. \**P*<0.05 (Student’s t-test, two-tailed)). **c,** FULVR (2) are independently derived fulvestrant-resistant MCF7 cells^59^. RT-qPCR was performed using RNAs prepared from FULVR cell lines and parental fulvestrant-sensitive cell lines. Heatmap shows the expression of TEADs and TEAD co-factors in FULVR cells. Gene expression values are represented as log2 (fold change) relative to the isogenic parental cell lines and normalized to GAPDH. **d,** Immunoblotting showing VGLL1 expression in different FULVR cell lines, accompanied by reductions in ER and PR levels. **e,** Normalised read counts from RNA-seq performed using MCF7 and isogenic LTED, TAMR and FULVR cells (2 bio-replicates samples)^13^. **f,** Immunoblotting of MCF7 cells following treatment with fulvestrant for 48 hours. **g,** RT-qPCR was carried out following treatment of cell lines with 100 nM fulvestrant for 48 hours. Gene expression data are shown as in **c**. **h,** RT-PCR was performed using RNA prepared from MCF7 cells treated with GDC0810, AZD9496 or RAD1901 for 48 hours. **i,** Immunoblotting of protein lysates prepared from MCF7 cells transfected with siESR1 or a control siRNA. **j,** Heatmap shows difference in expression of TEADs and TEAD co-factors using RT-qPCR of RNA prepared 48 hours following siESR1 transfection.

**Extended Data Figure 2.**
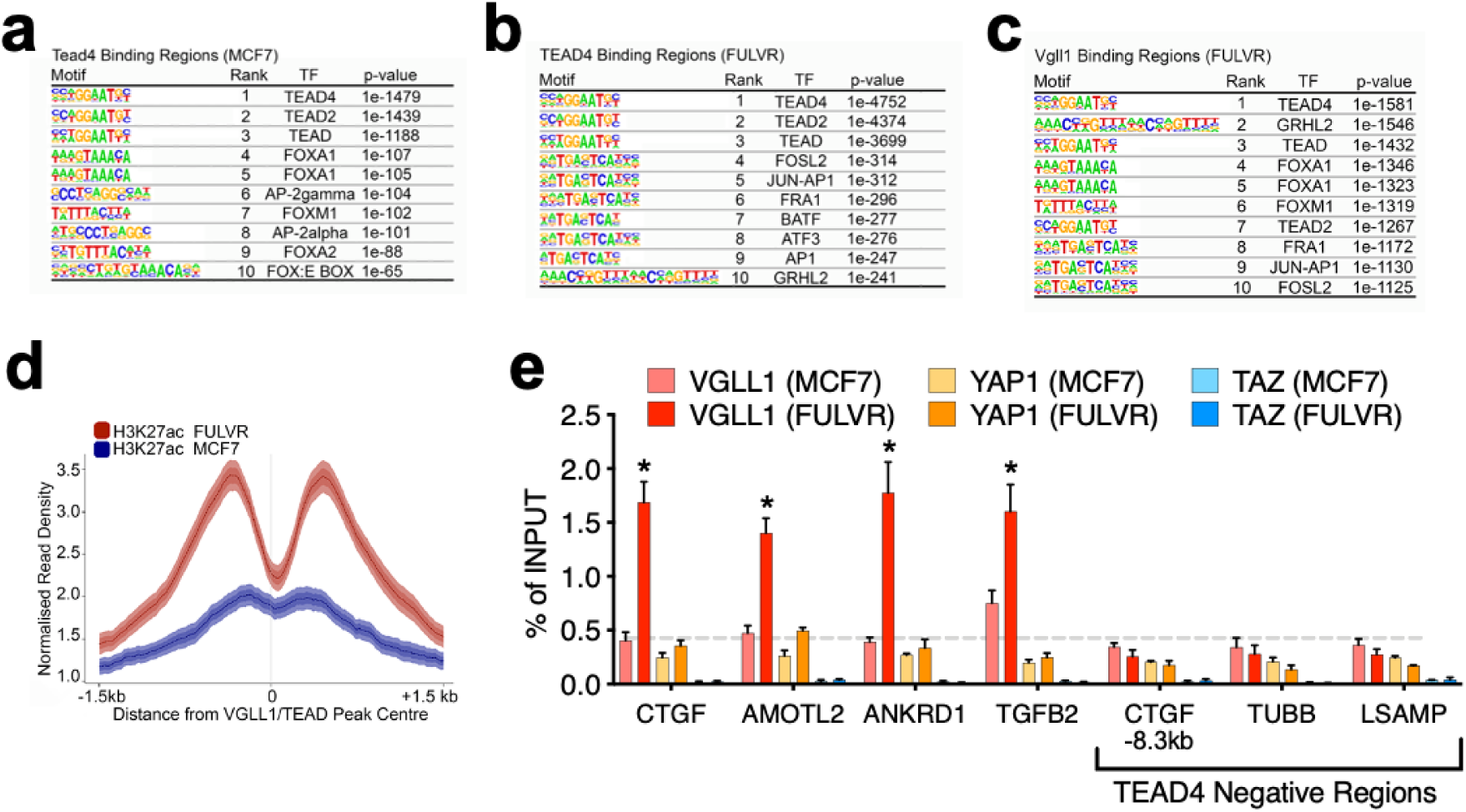
VGLL1 is recruited to gene regulatory regions in FULVR cells. **a-c,** *De novo* motif analysis of TEAD4 peaks in MCF7 cells and TEAD4 and VGLL1 peaks in FULVR cells. **d,** Normalized H3K27ac signal at TEAD4/VGLL1 co-bound regions from FULVR cells showing lower H3K27ac signal at these regions in MCF7 cells compared to FULVR cells. **e,** ChIP was performed using antibodies for VGLL1, YAP or TAZ, followed by qPCR for promoter and enhancer regions at TEAD binding regions, or regions that do not bind YAP/TAZ or TEAD (negative controls), selected from a previous report^53^. Binding enrichment was calculated as percentage of input. \**P*<0.05 (Student’s t-test, two-tailed).

**Extended Data Figure 3.**
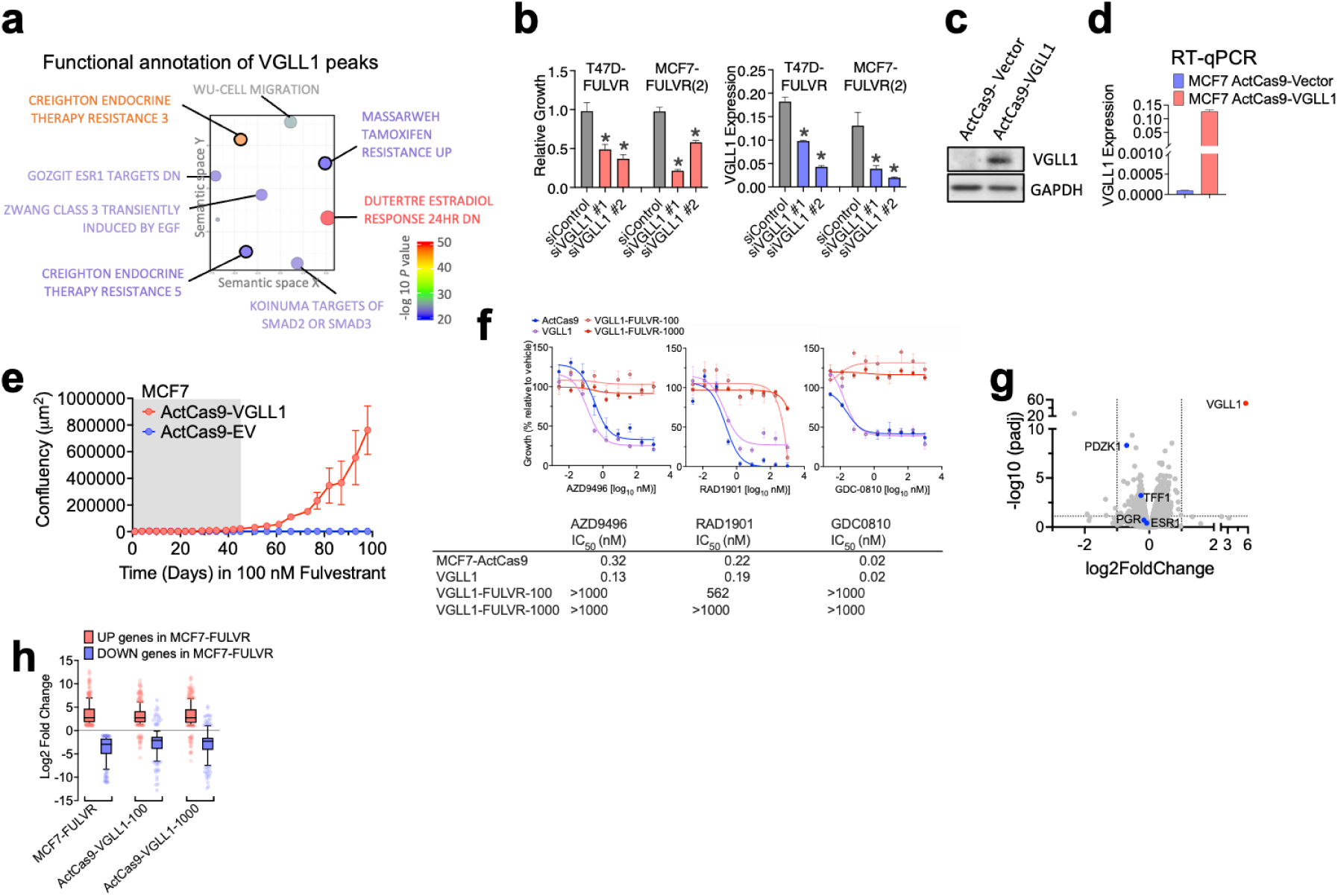
VGLL1 drives FULVR cell growth. **a,** Functional annotation of the VGLL1 ChIP-seq peaks in FULVR cells was performed using GREAT^60^. REVIGO^61^ was used to visualize the top nine most significant categories (see Supplementary Table 4 for the complete list). **b,** Left: Growth of T47D-FULVR cells and MCF7-FULVR(2) determined with the SRB assay five days after VGLL1 siRNA transfection. Growth was calculated relative to the non-targeting siRNA control (siControl). Data are mean and s.e.m of n=6. \**P* < 0.05 (Mann-Whitney test, two tailed). Right: VGLL1 gene expression after VGLL1 siRNA transfection in the indicated cell lines (*n*=3, \**p*<0.05 (Student’s t-test, two-tailed)). **c-d,** Immunoblotting (c) and RT-qPCR (d) of MCF7 ActCas9-VGLL1 and control cells (MCF7 ActCas9-Vector). **e,** Growth of MCF7 ActCas9-VGLL1 cells and control MCF7 ActCas9-Vector cells monitored using live cell imaging over a period of 100 days in the presence of 100 nM fulvestrant. The shaded area represents the part of this graph shown in Fig. 3c. **f,** Growth of the indicated MCF7-ActCas9 cells following five days of treatment with increasing doses of new generation SERDs (AZD9496, RAD1901, GDC-0810). Data are plotted as the mean ± s.e.m of n=6. **g,** RNA-seq data represented as a volcano plot for the comparison between MCF7 ActCas9-VGLL1 vs MCF7 ActCas9-Vector cells. **h,** Differential expression of DEGs up-regulated in FULVR cells relative to fulvestrant-sensitive MCF7 cells (log2FC > 2, p<0.05) and downregulated DEGs (log2FC < −2, p<0.05) was assessed in MCF7-ActCas9-VGLL1-FULVR-1 and in MCF7-ActCas9-VGLL1-FULVR-1 and the corresponding parental cells. The results indicate that expression of the FULVR DEGs is similarly altered in the VGLL1-over-expressing FULVR cells. Data are presented as box-and-whiskers plots (whiskers extend from the 5th to the 95th percentile; the box extends from the 25th to the 75th percentile; the line within the box represents the median).

**Extended data Fig. 4.**
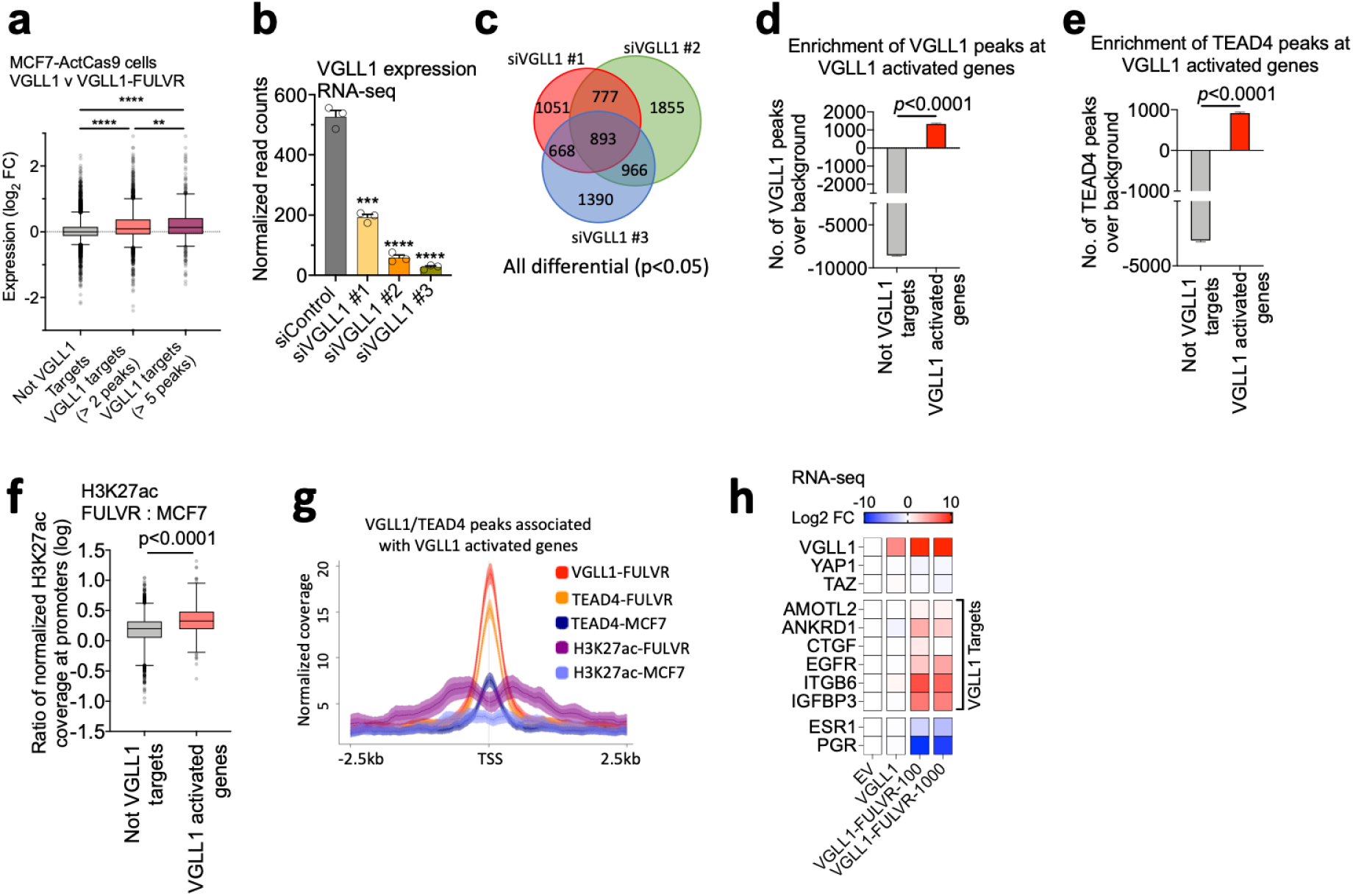
VGLL1-regulated genes are highly expressed in fulvestrant-resistant breast cancer cells. **a,** MCF7-ActCas9-VGLL1-FULVR cells show high expression of predicted VGLL1 targets. Genes with 2-4 and ≥5 VGLL1 peaks were considered as VGLL1 targets for this analysis. The y-axis shows the fold change in gene expression determined by RNA-seq between MCF7-ActCas9-VGLL1-FULVR-1 and the parental MCF7-ActCas9-VGLL1 cells. Data are presented as box-and-whiskers plots (whiskers extend from the 5th to the 95th percentile; the box extends from the 25th to the 75th percentile; the line within the box represents the median). \*\*\*\**P*<0.0001, \*\**P* = 0.0056 (Mann-Whitney test, two tailed). **b,** RNA-seq was carried out for 3 independent VGLL1 siRNAs, or a control siRNA (n=3 bioreplicates). Normalized read counts for VGLL1 are shown. **c,** Venn diagram showing all DEGs for the three VGLL1 siRNAs. **d,** VGLL1 binding is enriched at VGLL1 activated genes in FULVR cells. The y-axis shows the total number of VGLL1 peaks associated with all VGLL1-activated genes and not-VGLL1 targets over the mean expected value after generating random permutations of the VGLL1 peaks. *P*<0.0001 (Mann-Whitney test, two tailed). **e,** TEAD4 binding is enriched at VGLL1 activated genes in FULVR cells. **f,** Fold change in H3K27ac signal between FULVR versus MCF7 cells at the promoters of VGLL1 activated genes (*n* = 762) and not-VGLL1 targets (*n* = 8,932). Data are presented as box-and-whiskers plots, as in **a**. *P*<0.0001 (Mann-Whitney test, two tailed). **g,** Increased H3K27ac and TEAD4 recruitment in FULVR cells compared to MCF7 cells at regulatory elements associated with VGLL1 activated genes. Data are presented as average normalized ChIP-seq signal of VGLL1, TEAD4 and H3K27ac in FULVR cells together with TEAD4 and H3K27ac signal in MCF7 cells, in a window of ± 2.5 kb centred on the VGLL1/TEAD4 peaks associated with VGLL1 activated genes in FULVR cells. **h,** Heatmap showing Log2 FC in expression of VGLL1-regulated genes from RNA-seq analysis of MCF7-ActCas9-VGLL1-FULVR cells, relative to the vector control (EV) cells shows increased expression of VGLL1 and VGLL1 target genes, with a concurrent downregulation of ER transcriptional activity as evidenced by reduction in *PGR* expression. No change in YAP and TAZ expression is observed.

**Extended data Fig. 5.**
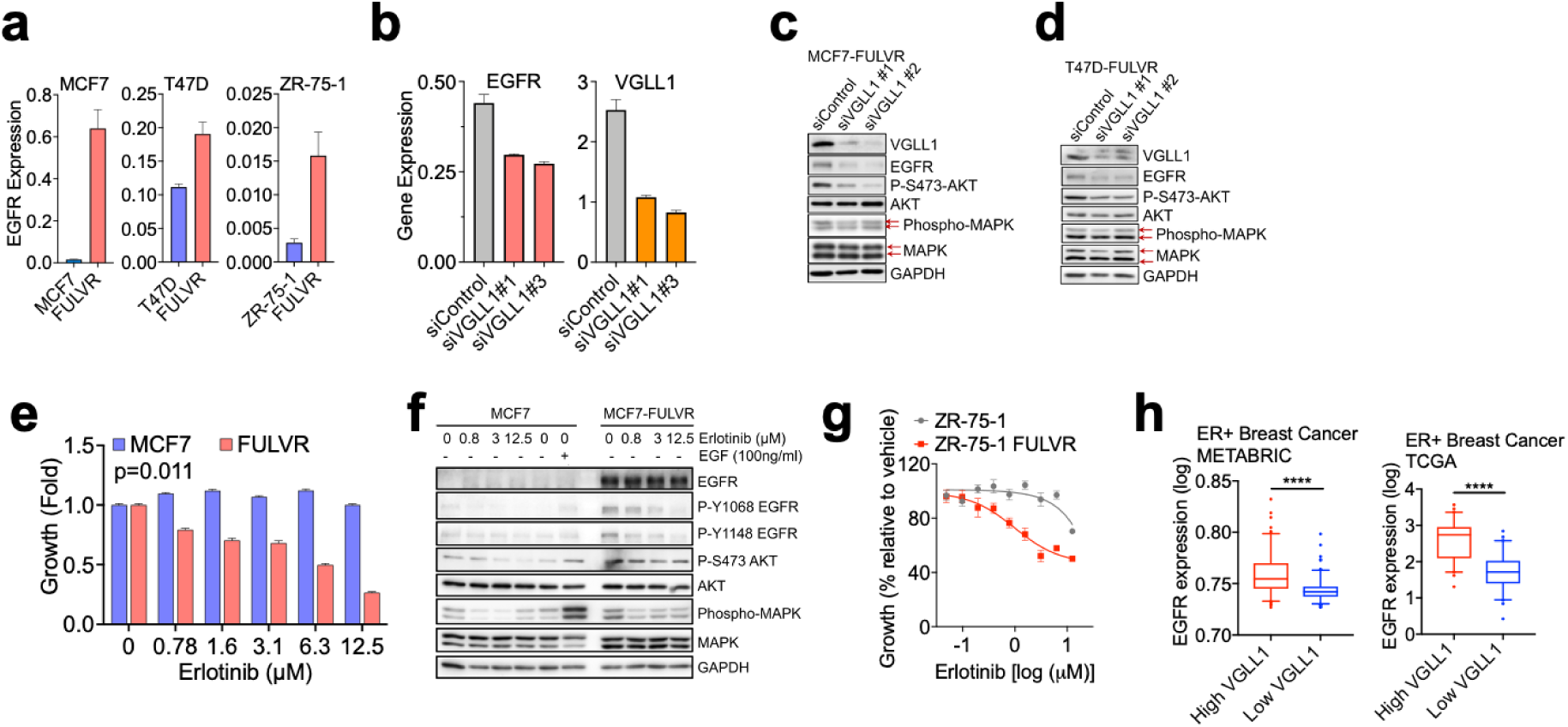
VGLL1 Induces EGFR expression in fulvestrant-resistant cells. **a,** RT-qPCR for EGFR showing EGFR upregulation in different fulvestrant resistant cell lines. **b,** RT-qPCR was undertaken using RNAs prepared from MCF7-ActCas9-VGLL1-FULVR cells transfected with two VGLL1 siRNAs. **c-d,** Immunoblotting of cell lysates prepared from MCF7-FULVR and T47D-FULVR cells 48 hours following transfection with VGLL1 siRNAs. **e,** MCF7 and MCF7-FULVR cells were treated with increasing concentrations of the EGFR inhibitor erlotinib for 5 days. Growth is shown as percentage relative to vehicle (n=6 replicates). **f,** Immunoblotting was carried out using cell lysates prepared from MCF7 and MCF7-FULVR cells treated with increasing concentrations of erlotinib for 24 hours. **g,** Growth of ZR-75-1 and the isogenic FULVR cells was assessed as in e. **h,** EGFR expression in ER+ breast cancer patients with high (top 10%) and low (bottom 10%) VGLL1 expression from TCGA and METABRIC datasets. ****P<2.2×10^-16^ (Mann-Whitney test, two tailed).

**Extended data Fig. 6.**
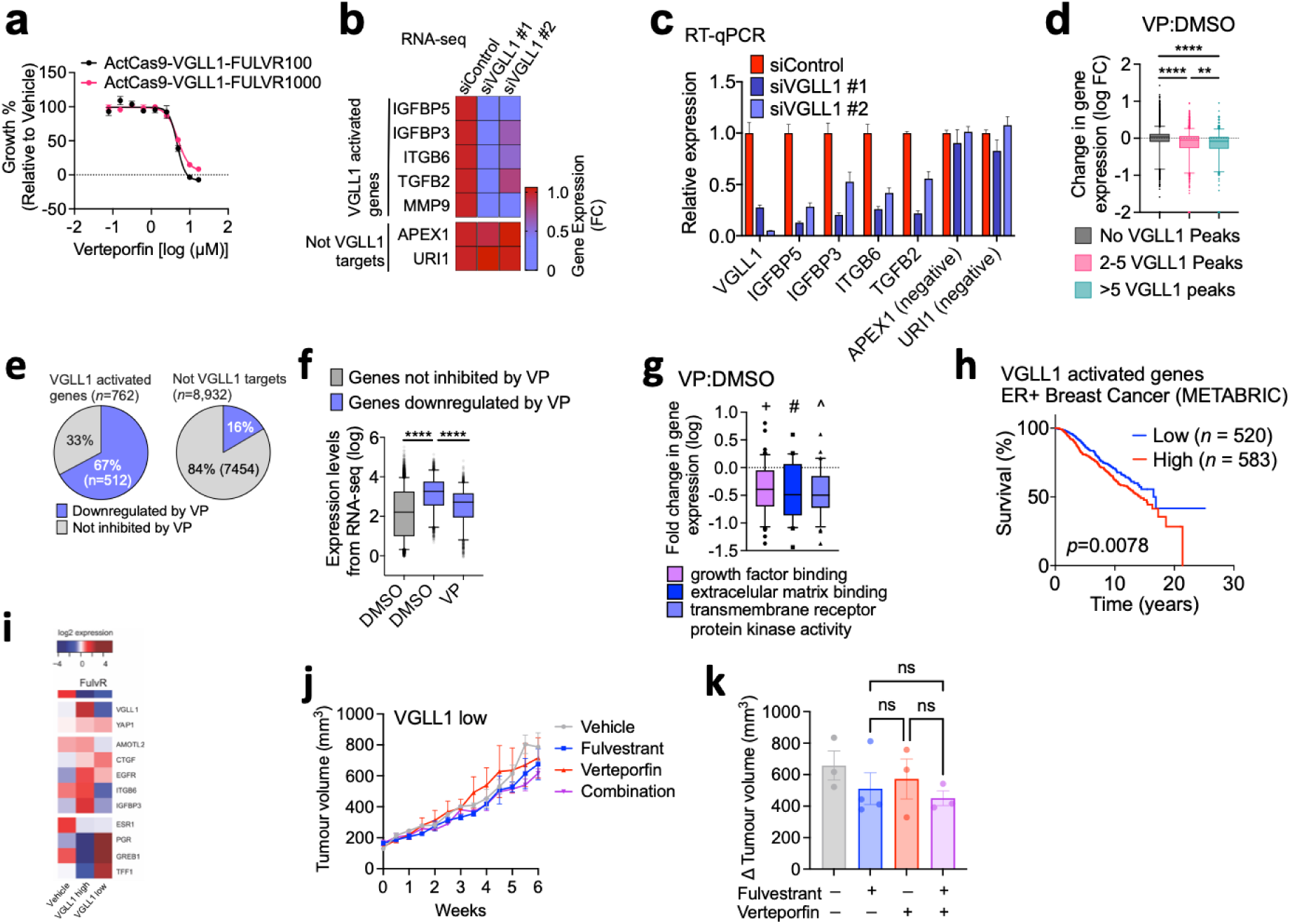
Verteporfin inhibits expression of VGLL1-regulated genes in fulvestrant-resistant breast cancer cells. **a,** Growth of MCF7 ActCas9-VGLL1-FULVR cells is inhibited by VP (n=6 replicates). **b,** Heatmap showing the expression of representative VGLL1 activated genes from RNA-seq of siVGLL1-treated MCF7-FULVR cells. **c,** RT-qPCR for *VGLL1*, representative VGLL1 activated genes, and not-VGLL1 targets (APEX1 and URI1) in MCF7-FULVR cells after siVGLL1 treatment. Expression values are represented as fold change relative to siControl following normalization to GAPDH levels. **d,** Genes associated with VGLL1 peaks are more prominently downregulated by VP compared to genes without VGLL1-associated peaks. \*\*\*\**P*<0.0001 (Mann-Whitney test, two tailed); ** *P* < 0.01 (Mann-Whitney test, two tailed). **e,** VGLL1 activated genes display high sensitivity to VP downregulation. Pie charts show the proportion of VGLL1-activated genes and not-VGLL1 targets genes downregulated by VP. **f,** Box-and-whiskers plot showing that genes downregulated by VP in FULVR cells (*adjusted P* < 0.01, log2(fold change) < −1) display higher expression levels than genes not inhibited by VP (*adjusted P* > 0.05, log2(fold change) > −1). \*\*\*\**P*<0.0001 (Mann-Whitney test, two tailed). **g,** Genes downregulated by VP are associated with growth functional categories enriched in the VGLL1 activated genes. Fold change in gene expression from RNA-seq of VP versus DMSO treated cells for genes included in the following GO terms: Growth factor binding (GO:0019838), extracellular matrix binding (GO:0050840) and transmembrane receptor protein kinase activity (GO:0019199), showing that genes included in these categories are significantly downregulated by VP. + *P_adj_* = 2.39×10^-25^, # *P_adj_* = 2.76×10^-9^, ^ *P_adj_* = 1.34×10^-22^ (Adjusted *p*-*values* from the pathway enrichment analysis of VP downregulated genes, see also Supplementary Table 6). **h,** Analysis of RNA-seq (n=2) data KCC_P_3837 PDX model following passaging in the presence of vehicle or fulvestrant, in FulvR clones 1 and 2. **i,** Tumour growth curves for mice bearing the KCC_P_3837 FulvR clone 1 (VGLL1-low) PDX model, treated as shown. **j,** Mean tumour volumes at 6 weeks with dots showing sizes of the individual tumours. ns = not significant in two-tailed unpaired t-test.

